# Phosphatase-Regulated Recruitment of the Spindle and Kinetochore Associated (SKA) Complex to Kinetochores

**DOI:** 10.1101/128827

**Authors:** Sushama Sivakumar, Gary J. Gorbsky

**Affiliations:** Cell Cycle and Cancer Biology Research Program, Oklahoma Medical Research Foundation, Oklahoma City, OK 77104, USA.

**Author notes:** Current Address: Department of Pharmacology, University of Texas Southwestern Medical Center, Dallas, TX 75390, USA. **Abbreviations** Ska: Spindle and Kinetochore-Associated PP1: Protein Phosphatase 1 PP2A: Protein Phosphatase 2A APC/C: Anaphase-Promoting Complex/Cyclosome.

**Keywords:** Mitosis, Cell Cycle, Anaphase-Promoting Complex, Spindle Checkpoint

## Abstract

Kinetochores move chromosomes on dynamic spindle microtubules and regulate cell cycle progression by signaling the spindle checkpoint. The Spindle and Kinetochore-Associated (Ska) Complex, a hexamer composed of two copies of Ska1, Ska2 and Ska3, participates in both roles. The mitotic kinases, Cdk1, Aurora B, Plk1, Mps1 and Bub1 play key, overlapping tasks in regulating chromosome movement and checkpoint signaling. However, roles for the phosphatases that oppose these kinases are more poorly defined. Recently, we showed that Ska1 is important for recruiting protein phosphatase 1 (PP1) to kinetochores. Here we show that PP1 and protein phosphatase 2A (PP2A) both promote accumulation of Ska at kinetochores. Depletion of PP1 or PP2A by siRNA reduces Ska binding at kinetochores, impairs alignment of chromosomes to the spindle midplane, and causes metaphase delay or arrest, phenotypes also seen after depletion of Ska. Tethering of PP1 to the kinetochore protein Nuf2 promotes Ska recruitment to kinetochores, and reduces mitotic defects seen after Ska depletion. We propose that kinetochore-associated phosphatases generate a positive feedback cycle to reinforce Ska complex accumulation and function at kinetochores.

**SUMMARY STATEMENT:** Phosphatases reinforce recruitment of the Ska complex at kinetochores to stabilize microtubule attachment and oppose spindle checkpoint signaling.

## INTRODUCTION

At mitotic entry, protein kinases phosphorylate numerous substrates to trigger important events such as chromosome condensation, nuclear envelope breakdown, mitotic spindle assembly, and proper kinetochore-microtubule interactions (Bollen et al., 2009; Wurzenberger and Gerlich, 2011). Throughout mitosis and particularly at mitotic exit, these phosphorylations are opposed by phosphatases. In mammalian cells, PP1 and PP2A, in association with specific regulatory and targeting subunits, are thought to dephosphorylate many of the substrates targeted by mitotic kinases (Bollen et al., 2009).

The three proteins of the Ska complex, Ska1, Ska2, and Ska3, exist as an obligate hexamer containing two copies of each (Gaitanos et al., 2009; Welburn et al., 2010). Depletion of any of the Ska proteins causes degradation of its partners, impaired chromosome alignment, and metaphase delay or arrest (Daum et al., 2009; Gaitanos et al., 2009; Hanisch et al., 2006; Redli et al., 2016; Schmidt et al., 2012; Sivakumar et al., 2014; Theis et al., 2009; Welburn et al., 2009). Both Ska1 and Ska3 bind to microtubules and promote chromosome movement (Abad et al., 2014; Abad et al., 2016; Schmidt et al., 2012). Ska1 also recruits protein phosphatase 1 (PP1) to kinetochores (Sivakumar et al., 2016), and the Ska complex promotes binding of the Anaphase-Promoting Complex/Cyclosome (APC/C) to mitotic chromosomes (Sivakumar et al., 2014). Phosphorylation of Ska by Aurora B kinase inhibits its binding to kinetochores, but paradoxically, a recent study reported that Ska also promotes Aurora B accumulation on kinetochores and increases its kinase activity *in vivo* and *in vitro* (Chan et al., 2012; Redli et al., 2016).

Two isoforms of PP1 (PP1γ and PP1α) are concentrated at kinetochores and bind Knl1 and Ska1 (Liu et al., 2010; Sivakumar et al., 2016; Trinkle-Mulcahy et al., 2006; Trinkle-Mulcahy et al., 2003). Kinetochore-associated PP1 appears to play important roles in stabilizing kinetochore-microtubule attachments and opposing spindle checkpoint signaling (Liu et al., 2010; Pinsky et al., 2006; Sivakumar et al., 2016; Vanoosthuyse and Hardwick, 2009).

The PP2A holoenzyme is a hetero-trimer composed of a scaffolding A subunit, regulatory B subunit and catalytic C subunit (Janssens et al., 2008). The B subunits are classified into three sub-families termed B (PR55/B55), B’(PR61/B56) and B’’(PR72) (Bollen et al., 2009; Janssens et al., 2008). Plk1 phosphorylation of BubR1 recruits PP2A-B56 to kinetochores in prometaphase (Foley et al., 2011; Suijkerbuijk et al., 2012). At metaphase, PP2A-B56 levels diminish at kinetochores while PP1 increases, suggesting that kinetochore-microtubule interactions are stabilized by PP2A-B56 in prometaphase and by PP1 at metaphase. In agreement with this idea, depletion of PP2A shows stronger impairment of chromosome alignment compared to depletion of PP1 (Foley et al., 2011; Liu et al., 2010).

In this study, we show that PP1 and PP2A phosphatases promote Ska recruitment to kinetochores. These results corroborate and extend previous work (Redli et al., 2016). Forced targeting of PP1 to kinetochores partially rescues defects caused by Ska3 depletion. We propose a feedback mechanism in which the Ska complex recruits PP1 to kinetochores at metaphase which further recruits Ska to stabilize kinetochore-microtubule attachments and initiate anaphase.

## RESULTS AND DISCUSSION

### PHOSPHATASES PROMOTE ACCUMULATION OF SKA AT KINETOCHORES

We and others have shown that Ska binds to kinetochores at prometaphase and maximally accumulates there at metaphase (Chan et al., 2012; Redli et al., 2016; Sivakumar et al., 2014). Inhibition of Aurora B kinase increased Ska accumulation on kinetochores lacking microtubule attachment (Chan et al., 2012). Correspondingly, expression of phosphomimetic mutants of Ska inhibited recruitment (Chan et al., 2012). These findings and recent data from Redli et al (2016) indicate phosphatases likely regulate Ska binding to kinetochores. PP1 and PP2A are the major phosphatases implicated in mitotic transitions. PP1, principally the PP1γ isoform, localizes to kinetochores and is implicated in spindle checkpoint inactivation (Liu et al., 2010; Trinkle-Mulcahy et al., 2003). PP2A also accumulates at kinetochores and plays a role in promoting kinetochore-microtubule attachment in prometaphase (Foley et al., 2011). To test the role of the phosphatases in Ska recruitment, we depleted PP1γ or PP2A Aα subunit (Figure S1A, S1B). We analyzed recruitment of Ska to kinetochores using immunofluorescence with antibody to Ska3. To normalize for the effect of microtubule binding, we performed these experiments in nocodazole-treated cells. Depletion of PP1 or PP2A phosphatase significantly decreased Ska3 accumulation at kinetochores (Figure 1A, 1B). Plk1 and BubR1 promote PP2A recruitment to kinetochores (Foley et al., 2011; Suijkerbuijk et al., 2012). Depletion of Plk1 or BubR1 with siRNA caused the expected reduction of PP2A at kinetochores in asynchronous cells (Figure S1C-E) and also resulted in lower levels of kinetochore-associated Ska3 in nocodazole (Figure 1C, 1D).

**Figure 1:**
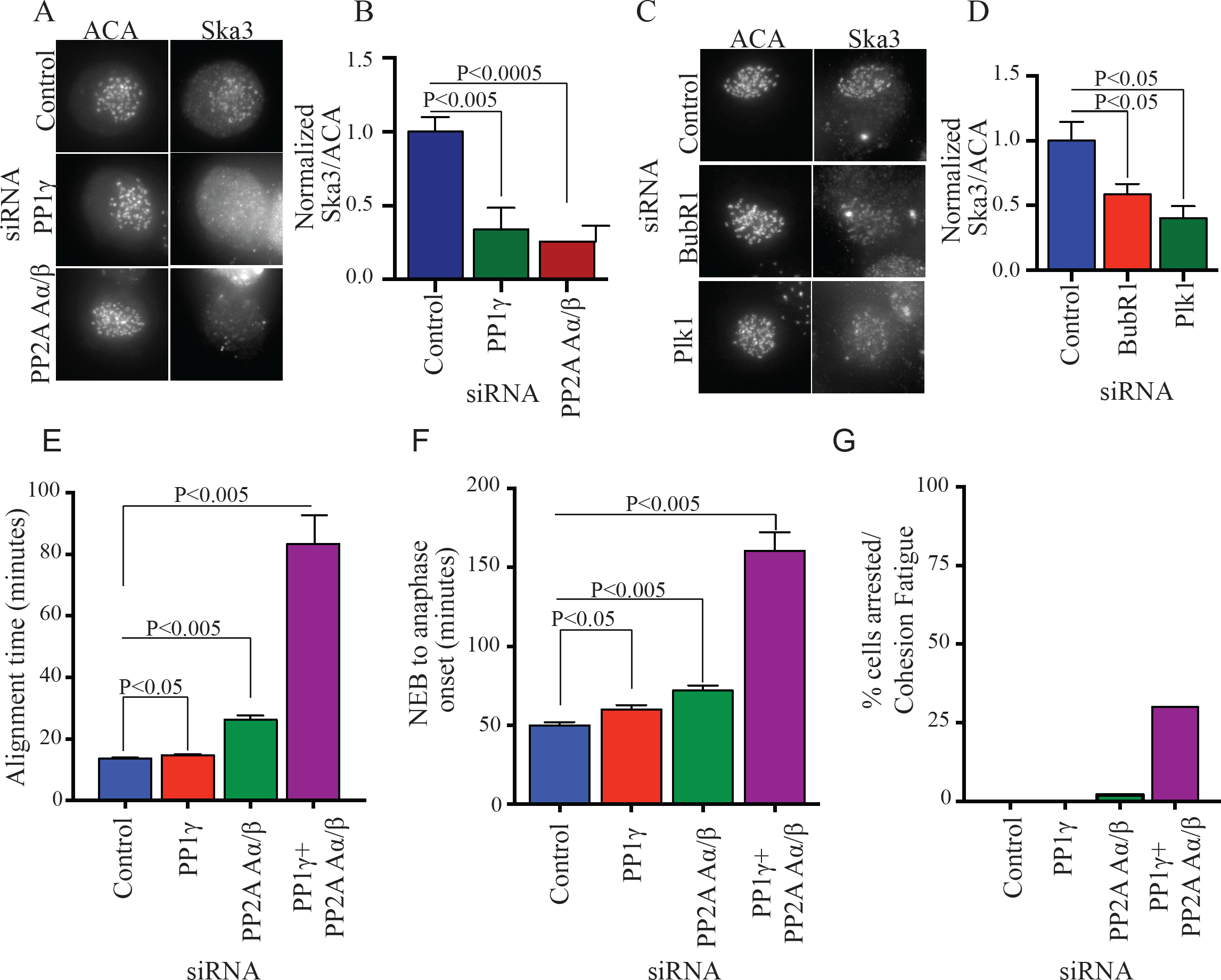
Phosphatases PP1 and PP2A promote Ska recruitment and normal progression through mitosis. A) HeLa cells grown on coverslips were transfected with control, PP1γ or PP2A Aα/β siRNA. 45h after transfection, cells were treated with 3.3μM nocodazole for 3h and then prepared for immunofluorescence. Ska3 at kinetochores was quantified. PP1γ or PP2A Aα/β depletion reduced Ska3 at kinetochore. B) Bar graph depicting mean fluorescence intensity of Ska3 at kinetochores normalized to anti-centromere antibody (ACA). C) HeLa cells grown on coverslips were transfected with control, Plk1 or BubR1 siRNA. 45h after transfection cells were treated for 3h with 3.3μM nocodazole and prepared for immunofluorescence. Ska3 at kinetochores was quantified. D) Plk1 or BubR1 depletion reduced Ska3 at kinetochores. E) HeLa H2B-GFP cells were transfected with PP1γ or PP2A Aα siRNA individually or in combination at 50nM final concentration. Approximately 30h post transfection, mitotic progression was followed by video microscopy. Bar graph depicts mean chromosome alignment times. PP1 depletion causes a slight delay in alignment. PP2A Aα depletion shows a stronger delay while combined depletion of PP1γ and PP2A shows the most robust delay. F) Bar graph depiction of time taken to initiate anaphase in individual cells. PP1γ or PP2A Aα depletions either individually or in combination delay mitotic progression. The combined depletions are more penetrant suggesting a compensatory or redundant role for the phosphatases when depleted individually. G) Bar graph showing percentage of cells siRNA-treated cells arrested in metaphase and undergoing cohesion fatigue after depleting PP1γ, PP2A Aα/β or both.

Since PP1 and PP2A promote Ska kinetochore binding we tested if depletion of the phosphatases in our hands generated phenotypes similar to Ska depletion. Depletion of PP1γ caused small delays in chromosome alignment (Figure 1E). As reported previously (Foley et al., 2011; Tang et al., 2006), depletion of PP2A Aα resulted in delayed chromosome alignment (Figure 1E). Combined depletions of both PP1γ and PP2A Aα/β delayed chromosome alignment yet more (Figure 1E).

Depletion of PP1_γ_ by RNAi delayed the metaphase-anaphase transition modestly, confirming its previously demonstrated role in opposing spindle checkpoint signaling (Liu et al., 2010). Depletion of PP2A Aα/β also delayed the metaphase-anaphase transition, but co-depletion of PP1_γ_ and PP2A Aα/β showed the strongest delay, consistent with both PP1 and PP2A promoting the metaphase-anaphase transition (Figure 1F) (Grallert et al., 2015; Lee et al., 2017).

As a second approach to test the roles of phosphatases and kinases in Ska kinetochore recruitment we used small molecule kinase and phosphatase inhibitors to examine effects on Ska recruitment to kinetochores. HeLa cells were treated in the presence of nocodazole to depolymerize microtubules and the proteasome inhibitor MG132 to block mitotic exit. As reported previously, inhibition of Aurora kinase increased recruitment of Ska to kinetochores (Chan et al., 2012). However, in contrast to previous findings we found that treatment of cells with an inhibitor of Mps1 also increased Ska recruitment (Figure 2A, 2B) (Chan et al., 2012). This finding is consistent with roles for Mps1 in recruitment of Aurora B and BubR1 to kinetochores (Krenn et al., 2014; Overlack et al., 2015; van der Waal et al., 2012; Zhang et al., 2014). As also reported in a recent study (Redli et al., 2016), we found that treatment of cells with the phosphatase inhibitor, okadaic acid, decreased Ska accumulation at kinetochores (Figure 2A, 2B), consistent with results from RNAi depletion of phosphatases described above. Previously we found that Ska complex promotes APC/C accumulation on mitotic chromosomes (Sivakumar et al., 2014). We tested if kinase or phosphatase inhibitors that affected kinetochore concentration of Ska had similar effects on chromosome-bound APC/C. Indeed we found that inhibitors of Aurora B or Mps1 increased APC/C on chromosomes while the phosphatase inhibitor, okadaic acid, decreased APC/C on chromosomes (Figure 2C, 2D).

**Figure 2:**
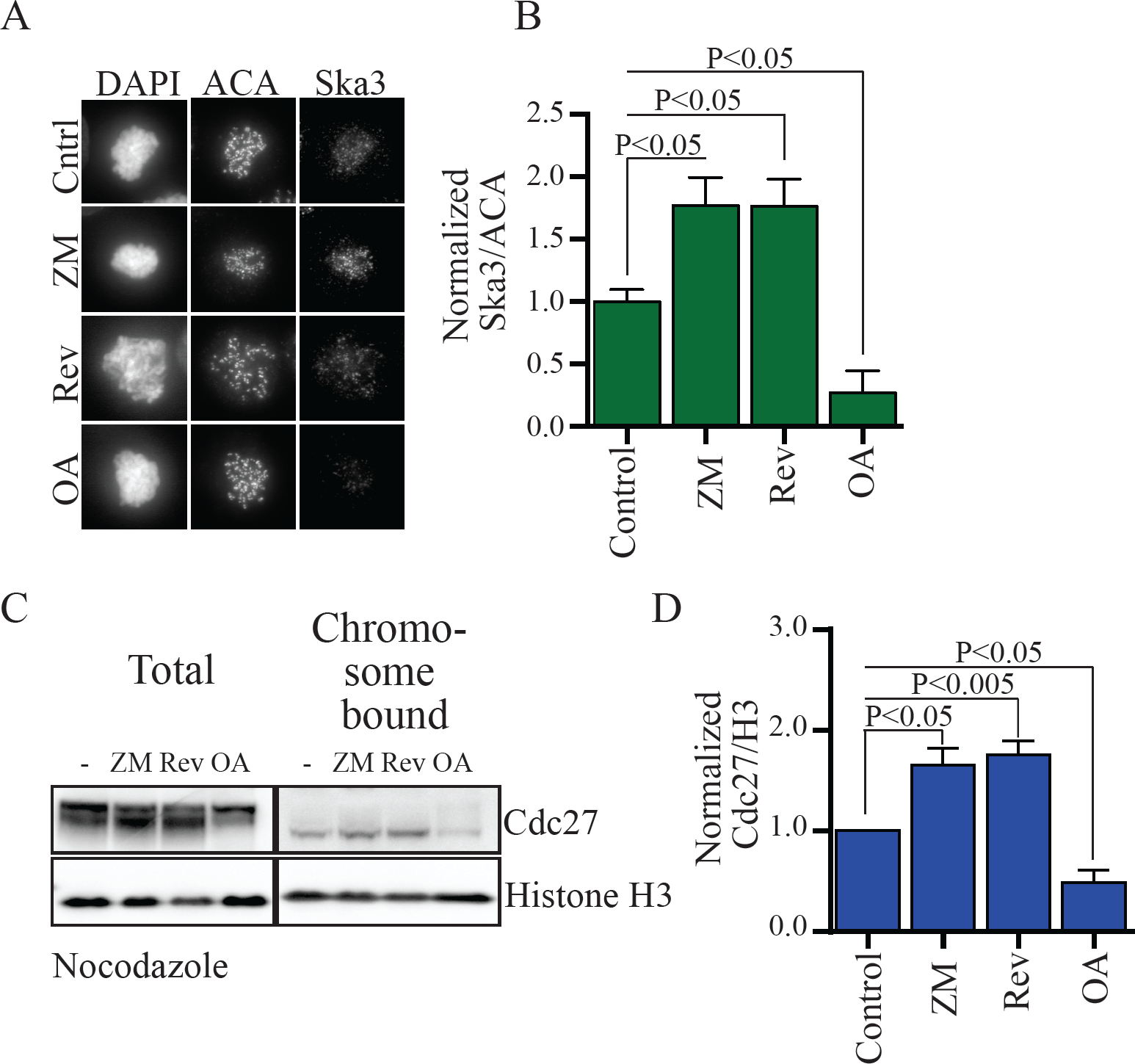
Kinase and phosphatase inhibitors regulate Ska recruitment to kinetochores and APC/C recruitment to mitotic chromosomes. A) HeLa cells grown on glass coverslips were treated with 3.3µM nocodazole for 2h. Coverslips were individually treated with DMSO (control), 1μM reversine (Mps1 inhibitor), 25µM ZM447439 (ZM-Aurora B inhibitor) or 500nM Okadaic acid (OA- phosphatase inhibitor) for an additional 1h. Cells were pre-extracted, fixed and labeled with anti-Ska3 antibody and ACA for immunofluorescence and quantified. B) Aurora B or Mps1 inhibition increases Ska3 at kinetochores while phosphatase inhibition decreased Ska3 at kinetochores. C) HeLa cells treated as in A were collected and mitotic chromosomes were isolated to analyze amount of APC/C associated. D) Blot quantification shows that Cdc27 levels increase in chromosomes treated with Aurora B inhibitor or Mps1 inhibitor and decrease in OA treated cells.

### MICROTUBULE ATTACHMENT AND PP1 PROMOTE SKA3 KINETOCHORE RECRUITMENT

We previously demonstrated that an important function of the C-terminus of Ska1 is to recruit PP1 to kinetochores (Sivakumar et al., 2016). In those experiments, Ska2, Ska3 and the N terminus of Ska1 and their potential separate functions were retained. We constructed a fusion of the outer kinetochore protein, Nuf2, to PP1. Nuf2 is a component of the Ndc80 complex, which is implicated in recruiting Ska to kinetochores (Chan et al., 2012; Zhang et al., 2012). We reasoned that the Nuf2PP1 fusion would target PP1 in close proximity to the normal Ska-recruited PP1. Expression of the Nuf2PP1 fusion induced a significant increase in kinetochore-associated PP1 both in cells arrested in a prometaphase-like state in nocodazole and at metaphase in MG132 (Figure 3A, 3B, 3C) even though the Nuf2PPI fusion was expressed at lower levels than unfused Nuf2 or PP1 in controls (Figure S2A). We then analyzed Ska recruitment. Interestingly, in nocodazole-treated cells, Nuf2PP1 had little effect on Ska accumulation at kinetochores (Figure 3D, S2B). However, in MG132-treated cells at metaphase, Nuf2PP1 caused a 50% increase in Ska accumulation at kinetochores (Figure 3E, 3F).

**Figure 3:**
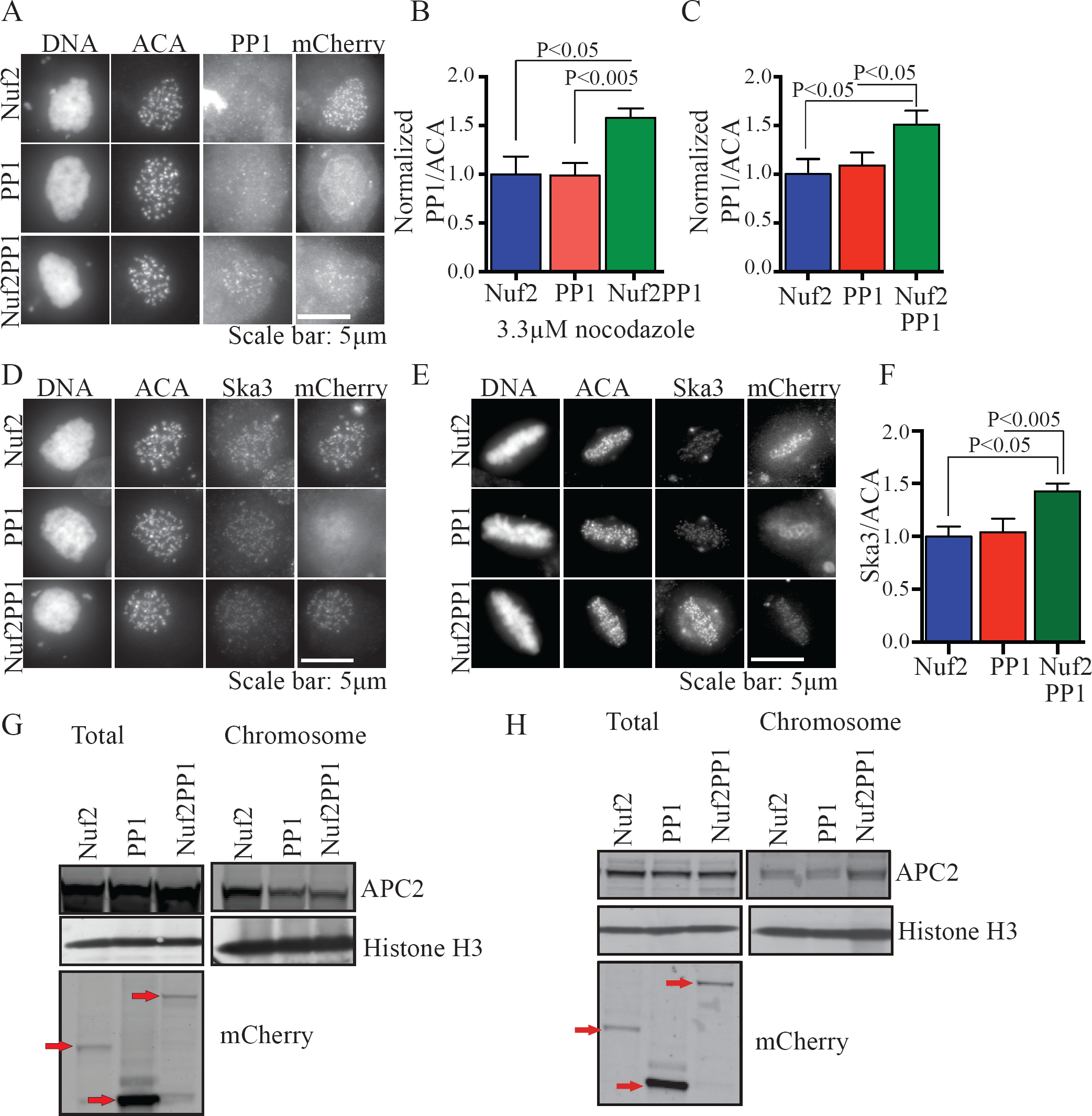
Expression of a Nuf2PP1 fusion in the presence of intact spindle microtubules promotes Ska recruitment to kinetochores and APC/C recruitment to chromosomes. A) HeLa cells grown on coverslips were transfected with Nuf2-mCherry, PP1-mCherry or Nuf2PP1-mCherry. 36h after transfection 3.3μM nocodazole was added to cells for 3h and immunofluorescence was done. PP1 localization to kinetochores was quantified. B) Nuf2PP1 expression increased PP1 at kinetochores by 50%. C) HeLa cells transfected with Nuf2-mCherry, PP1-mCherry or Nuf2PP1-mCherry were released from 330nM nocodazole into MG132 for 2h. Immunofluorescence was done and PP1 at kinetochores was measured. Nuf2PP1 expression increases PP1 at kinetochores in MG132 treated cells. D) HeLa cells were treated as in A and immunofluorescence was done. Amount of Ska3 at kinetochores was quantified. In cells treated with nocodazole, Ska3 at kinetochores was not different in cells expressing Nuf2-mCherry, PP1-mCherry, or Nuf2PP1-mCherry. E) HeLa cells were treated as in C and Ska3 at kinetochores was measured. F) Ska3 accumulation at kinetochores increases in MG132-treated cells expressing Nuf2PP1-mCherry. G) Cells were treated as in A and then used to prepare mitotic chromosomes. Chromosome bound APC/C is similar in cells expressing Nuf2-mCherry, PP1-mCherry, or Nuf2PP1-mCherry. H) Cells were treated as in C then used to prepare mitotic chromosomes. Chromosome-bound APC/C increases in MG132-arrested mitotic cells expressing Nuf2PP1-mCherry.

Previously we showed that Ska promotes binding of the APC/C to mitotic chromosomes (Sivakumar et al., 2014). Similar to the findings described above regarding Ska recruitment to kinetochores, we found that expression of Nuf2PP1 did not impact chromosome-bound APC/C in nocodazole-treated cells, but did increase it in cells arrested at metaphase with MG132 (Figure 3G, 3H, S2C).

### EXPRESSION OF NUF2PP1 FUSION PARTIALLY RESCUES MITOTIC DEFECTS IN SKA3-DEPLETED CELLS

Depletion of any Ska component results in delayed alignment followed by a robust mitotic delay or arrest at metaphase (Daum et al., 2009; Redli et al., 2016; Schmidt et al., 2012; Sivakumar et al., 2014). The metaphase delay often results in cohesion fatigue, asynchronous separation of chromatids without mitotic exit (Daum et al., 2011; Sivakumar et al., 2014; Stevens et al., 2011). We tested if Nuf2PP1 expression rescued mitotic defects in Ska-depleted cells. In control cells, expression of Nuf2, PP1 or Nuf2PP1 did not alter mitosis (Figure S2D). In Ska3-depleted cells, expression of the Nuf2PP1, but not Nuf2 or PP1 alone, improved alignment (Figure 4A, 4B). In addition, while 63% of Ska-depleted cells expressing Nuf2 or PP1 arrested at metaphase and underwent cohesion fatigue, only 22% of cells expressing the Nuf2PP1 fusion did so (Figure 4C). However, the rescue was incomplete, most Ska3-depleted cells expressing the Nuf2PP1 fusion still exhibited a significant delay before entering anaphase (Figure 4A). There are potential technical reasons why the rescues were incomplete. Both PP1 and Ska have been shown to exhibit rapid turnover at kinetochores (Raaijmakers et al., 2009; Trinkle-Mulcahy et al., 2001). In contrast the Ndc80 complex, of which Nuf2 is a component, is stable (Hori et al., 2003). Thus, Nuf2PP1 likely fails to exhibit the normal dynamics of PP1 and this limitation possibly prevents complete rescue of Ska depletion phenotypes.

**Figure 4:**
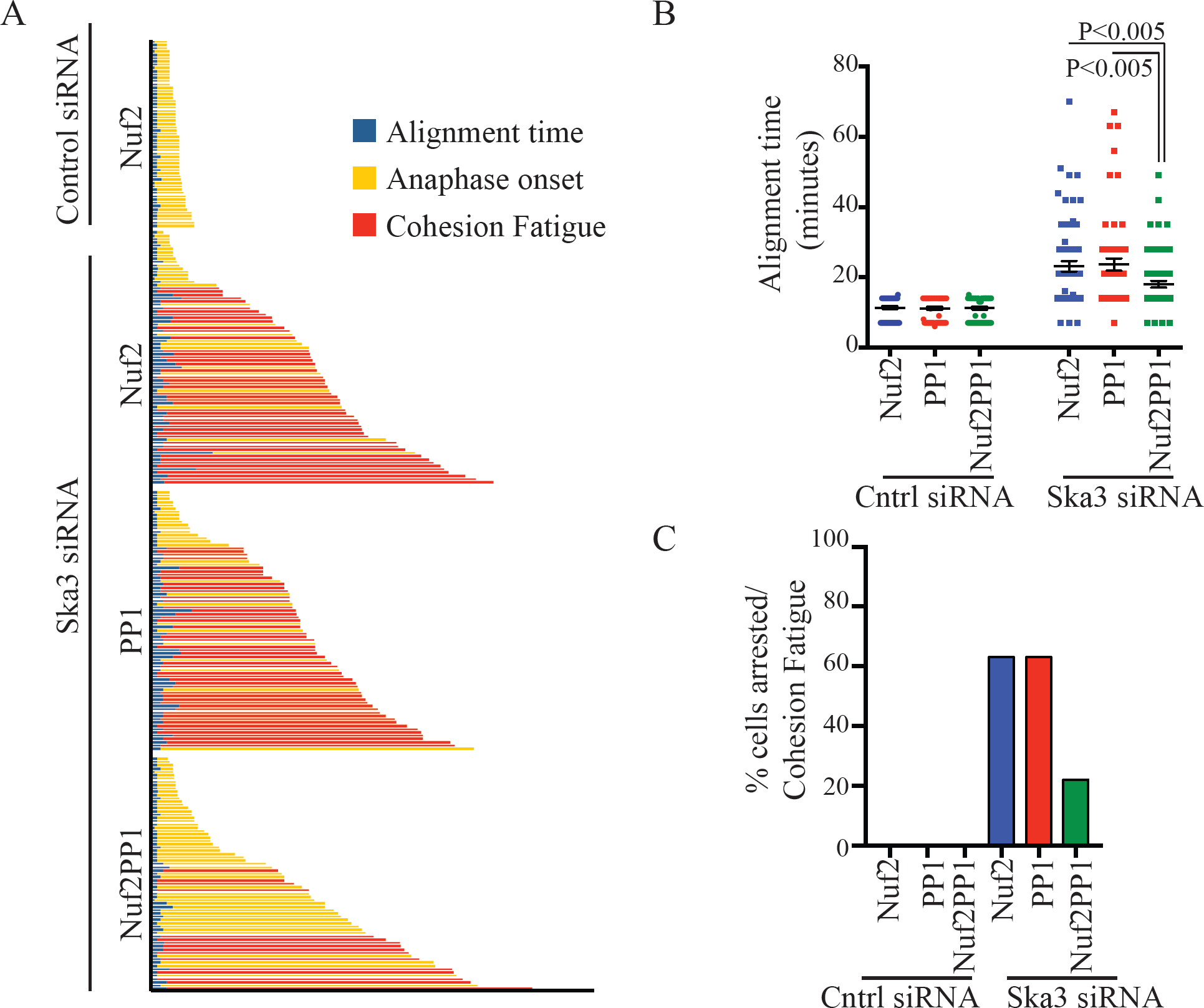
Expression of Nuf2PP1 fusion partially rescues mitotic defects caused by Ska depletion. A) Individual fates for HeLa H2B GFP cells transfected with Nuf2-mCherry, PP1-mCherry or Nuf2PP1-mCherry. 12h after transfection, cells were treated with control or Ska3 siRNA. Time-lapse imaging was done 24h after siRNA transfection. The intervals to align chromosomes, initiate anaphase or undergo metaphase arrest/cohesion fatigue were measured. B) Scatter plot depicts time taken to align chromosomes in control or Ska3-depleted Nuf2-mCherry, PP1-mCherry or Nuf2PP1-mCherry expressing cells. Expression of Nuf2PP1-mCherry partially rescued alignment in Ska3-depleted cells. C) Bar graph showing percentage of cells arrested at metaphase and undergoing cohesion fatigue in control or Ska3-depleted cells expressing Nuf2-mCherry, PP1-mCherry or Nuf2PP1-mCherry. Expression of Nuf2PP1 promotes anaphase onset in Ska3 depleted cells compared to Nuf2 or PP1 expressing cells.

## CONCLUSIONS

Here we show that PP1 and PP2A are required for full accumulation of Ska on kinetochores. Though cells depleted of PP1_γ_ and PP2A Aα/β show alignment defects and metaphase delays, they rarely exhibit the strong phenotypes characteristic of Ska depletions (Figure 1G). There are possible technical explanations for this observation. PP1 and PP2A proteins may be long-lived or be sufficient at low protein concentration. Other mitotic phosphatases may also compensate (Posch et al., 2010; Wurzenberger et al., 2012).

If PP1 recruitment were the primary function of Ska, expression of Nuf2PP1 in Ska-depleted cells might be expected to completely alleviate phenotypes caused by reduction of Ska levels. We found only partial rescue. Most likely, PP1 recruitment, a function of the C terminus of Ska1, is only one function of the full Ska complex. Additional roles such as microtubule tracking may also promote alignment and timely metaphase-anaphase transition (Abad et al., 2014; Abad et al., 2016; Chan et al., 2012; Gaitanos et al., 2009; Raaijmakers et al., 2009; Redli et al., 2016; Schmidt et al., 2012; Welburn et al., 2009).

PP1 has been shown to be integral in opposing spindle checkpoint signaling and is required to promote normal anaphase onset (Liu et al., 2010; Sivakumar et al., 2016). PP1 levels at kinetochores increase from prometaphase to metaphase. We speculate that low levels of PP1 and Ska at kinetochores and APC/C on chromosomes in prometaphase prevent premature anaphase onset and mitotic exit. Then, at metaphase Ska binding to microtubules increases its kinetochore concentration creating a positive feedback loop whereby PP1 accumulation at kinetochores further increases Ska leading to accumulation of kinetochore PP1 and chromosome APC/C. This feedback strengthens microtubule attachment, opposes spindle checkpoint signaling and promotes the rapid and irreversible transition to anaphase and mitotic exit.

## MATERIALS AND METHODS

### CELL CULTURE

HeLa cells stably transfected with GFP fused to Histone 2B (HeLa H2B-GFP) was used in this study. HeLa cell lines were grown in culture flasks or chambered coverslips in DMEM-based media with 10% FBS supplemented with penicillin and streptomycin in 5% CO_2_ at 37°C. Cell lines were routinely tested for mycoplasma contamination and only used if not contaminated.

Transient transfection of siRNA was done using Lipofectamine RNAi reagent (Invitrogen) according to manufacturer’s instructions. siRNA against PP2A Aα/β (CACAGAGAAAUAAAGGUCU and ACAACGUCAAGAGUGAGAU), PP1_γ_ (CGAGUGACCGAUUAUGCUU and GUCUGAGGAGUAAGUGUAC) were obtained from Bioneer Inc. and these were used at 50-100nM final concentration. siRNA against Ska3 was obtained from Dharmacon and used at 50nM final concentration (Daum et al., 2009).

### PLASMIDS

Nuf2, PP1 or Nuf2PP1-mCherry plasmids were constructed by inserting respective cDNA in mCherry-N1 vector. mCherry tag was inserted upstream or downstream of transgene and similar results were obtained with proteins tagged in N or C termini.

### LIVE CELL IMAGING

HeLa H2B-GFP cells were grown in Nunc chambered coverslips (Thermo Sci. Inc.). To maintain appropriate pH levels and avoid evaporation during imaging, culture media was exchanged to Leibovitz’s L-15 medium supplemented with 10% FBS, penicillin, streptomycin and overlaid with mineral oil. Time-lapse fluorescence images were collected using 20X or 40X objectives and a Zeiss Axiovert 200M inverted microscope equipped with an objective heater, air curtain, a Hamamatsu ORCA-ER camera, and Metamorph software (MDS Analytical Technologies). Images were captured every 4 – 15 minutes for 12 – 24 hours. Time-lapse videos displaying the elapsed time between consecutive frames were assembled using Metamorph software. The mitotic interval was calculated from nuclear envelope breakdown (NEB) until anaphase onset/mitotic exit and depicted as bar graphs or scatter plots with mean and SEM. In scatter plots, each dot represents one cell; long horizontal line depicts mean and whiskers denote SEM. The unpaired Student T test in Graphpad Prism was used to assess statistical significance.

### IMMUNOFLUORESCENCE AND QUANTIFICATION

HeLa cells were grown on glass coverslips and treated as detailed in the figure legends. Cells were pre-extracted in PHEM/1% triton solution for 5 minutes before fixing with 1.5% paraformaldehyde/PHEM solution for 15 minutes. Coverslips were washed in MBST, blocked in 20% boiled goat or donkey serum, and incubated overnight with primary antibodies. Samples were then incubated with secondary antibodies for 1-2 h, stained with DNA dye, DAPI, and mounted using Vectashield (Vector Laboratories). The following primary antibodies were used: rabbit anti-Ska3 (Daum et al., 2009), ACA/CREST (Antibody Inc.), mouse anti-Bub1 (antibody 4B12 from Dr. Steven Taylor, University of Manchester), rabbit anti-Plk1 (Upstate), rabbit anti-BubR1 (gift from Dr. Todd Stukenberg), goat anti-PP1γ (Santa Cruz Inc.), goat anti-PP2A Aα/β (Santa Cruz Inc.). Secondary antibodies used were goat anti–rabbit, goat anti–mouse, donkey anti-goat antibodies conjugated to Cy3 or FITC or goat anti-human antibody conjugated to Cy3 or FITC (Jackson ImmunoResearch). The images were acquired using Zeiss Axioplan II microscope equipped with a 100X objective (N.A. 1.4), a Hamamatsu Orca 2 camera (Hamamatsu Photonics) and processed using MetaMorph software (Molecular Devices) and Coreldraw (Corel Corp.). Quantification of the immunofluorescence images was done as described previously (Daum et al., 2009; Sivakumar et al., 2014). 5-10 cells in each condition were quantified. The graphs depict average fluorescence value with SEM in each condition. Every experiment was repeated at least 3 times and representative images are shown.

### WESTERN BLOTTING

Whole cell HeLa cell extracts were prepared by lysis in APCB buffer (20mM Tris-Cl pH 7.7, 100mM KCl, 50mM Sucrose, 1mM MgCl2, 0.1mM CaCl2, 0.5% Triton X-100) containing protease inhibitor cocktail (Sigma Aldrich) and microcystin (400nM). For electrophoresis, sample loading buffer (Invitrogen) and dithiothreitol (DTT) to a final concentration of 50mM were added. Proteins were separated with a NuPAGE gel electophoresis system (Invitrogen), transferred to 0.45µm PVDF membrane (Immobilon PVDF, Millipore) via a Genie transfer apparatus (Idea Scientific). Membranes were blocked in 5% Non-Fat Dry Milk (NFDM) and 0.05% Tween 20 in Tris-buffered saline (TBS). Primary antibodies included rabbit anti-Ska3 antibody (Daum et al., 2009), mouse anti-β-Actin (Abcam), anti-Plk1 (Upstate), anti-BubR1 (gift from Todd Stukenberg). Membranes were washed in TBS/0.05% Tween 20 (TBST), and then incubated with secondary antibodies in 5% NFDM/TBST. Secondary antibodies include HRP goat anti-mouse or anti-rabbit antibodies (Jackson Immunoresearch). After washes, membranes were developed using West Pico Chemiluminescent reagent (Pierce) and imaged using a Kodak 4000M imaging station. Quantification of western blots was done as described previously (Sivakumar et al., 2014; Sivakumar et al., 2016). Every experiment was repeated at least 3 times.

### CHROMOSOME PREPARATION

HeLa cells were grown in 150mm plates and treated as indicated in figure legends. Mitotic cells were lysed with ELB buffer on ice (1X PHEM, 0.5% Triton X-100, 1mM DTT, 10% glycerol, protease inhibitors and 400nM microcystin) and centrifuged to separate cytoplasmic fractions from chromosome fractions. Chromosome fractions were further washed with ELB at least 3 times to remove cytoplasmic contamination and resuspended in 1/4^th^ volume ELB of cytoplasmic fractions. The protein concentration of cytoplasmic fractions was determined using the BCA protein assay kit (Pierce). The chromosomes were DNAse treated and resuspended in sample loading buffer (Invitrogen) and 50mM DTT. Samples were immunoblotted with antibodies to the APC/C components Cdc27 and APC2 and antibody to Histone H3.

## ACKNOWLEDGEMENTS

We thank Dr. Steven Taylor and Dr. Todd Stukenberg for providing antibodies. We thank Dr. Hongtao Yu for providing laboratory space and APC2 antibody. GJG was supported by RO1GM111731 from the National Institute of General Medical Sciences and by the McCasland Foundation.

## SUPPLEMENTAL FIGURE LEGENDS

**Figure S1:**
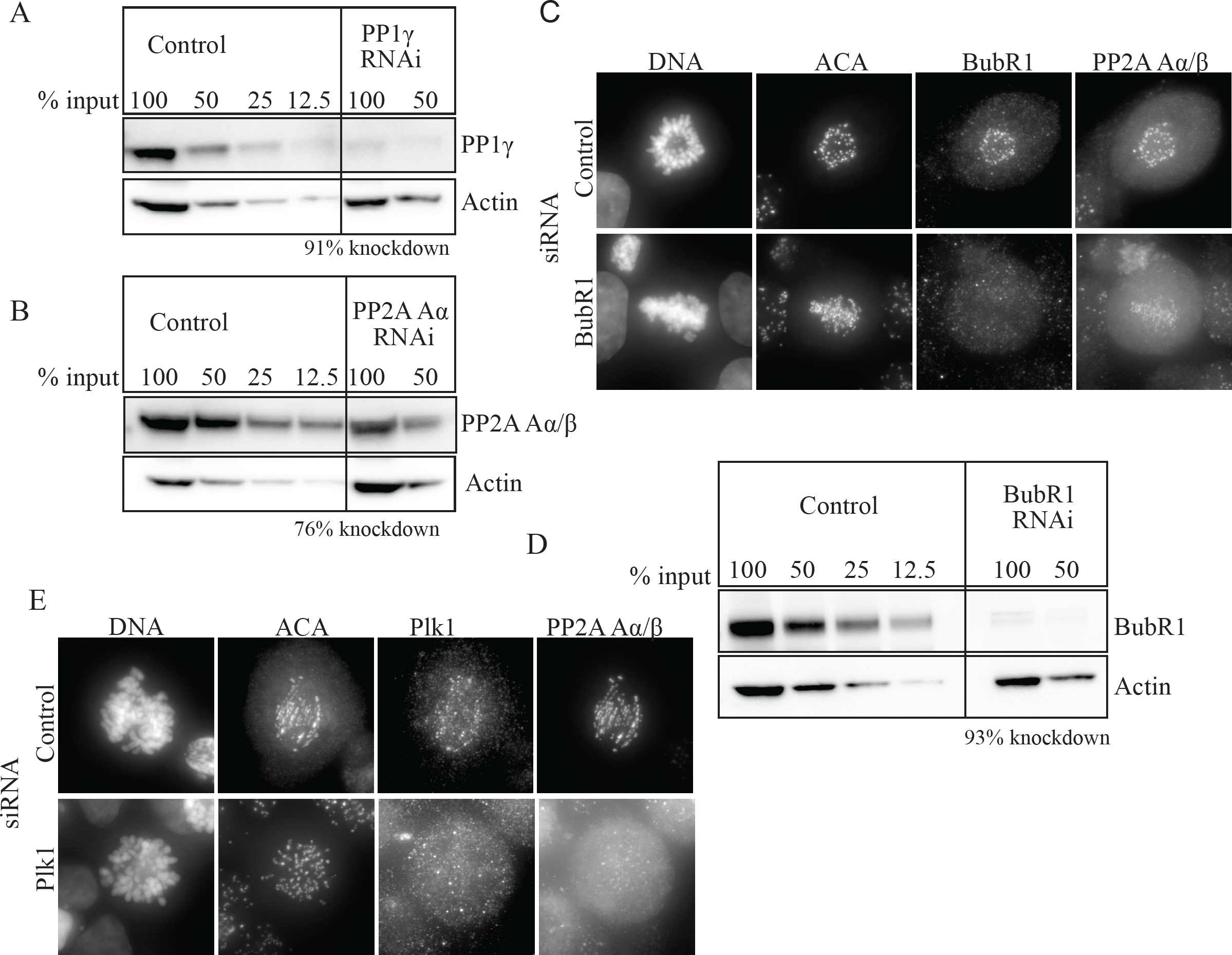
Depletions achieved by siRNA. A) HeLa cells transfected with PP1_γ_ siRNA were arrested in mitosis with 3.3μM nocodazole and collected. Western blotting of whole cell lysates indicated that PP1_γ_ was depleted by 91%. B) HeLa cells transfected with PP2A Aα/β siRNA were arrested in mitosis with 3.3μM nocodazole and collected. Western blotting of whole cell lysates indicated that PP2 Aα/β was depleted by 76%. C) HeLa cells transfected with control or BubR1 siRNA. Immunofluorescence was done and cells were stained for BubR1 and PP2A Aα/β. BubR1 depletion reduced BubR1 by 75 % and PP2A Aα/β by 40 % at kinetochores. D) HeLa cells transfected with BubR1 siRNA were arrested in mitosis with 3.3μM nocodazole and collected. Western blotting of whole cell lysates indicated that BubR1 was depleted by 93%. E) HeLa cells were transfected with control or Plk1 siRNA. Cells were immunolabeled with Plk1 and PP2A Aα/β. PP2A Aα/β was reduced by 50% in Plk1 depleted cells.

**Figure S2:**
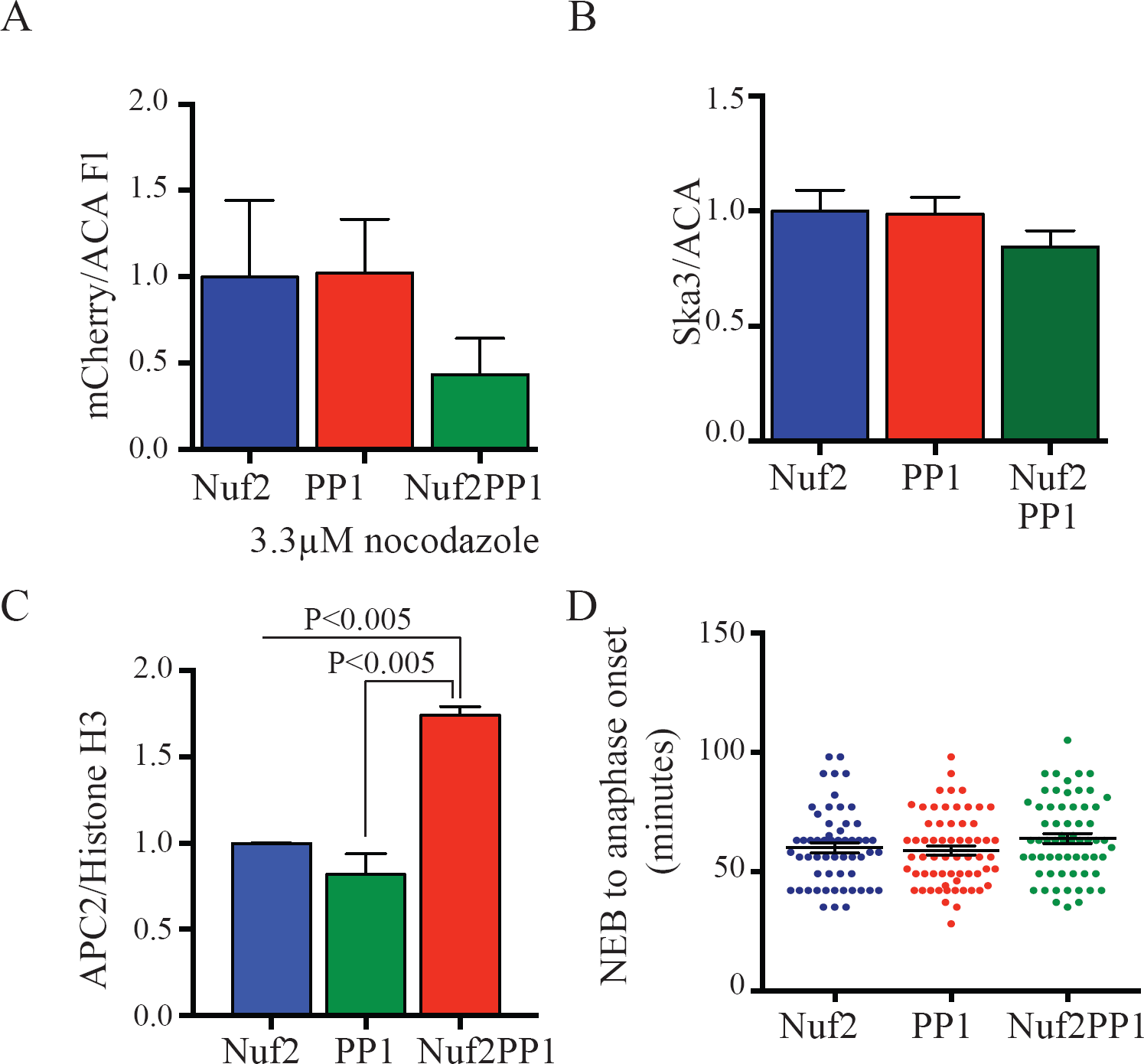
Expression of Nuf2PP1 does not increase kinetochore-associated Ska in nocodazole-arrested cells but does increase APC/C on chromosomes in MG132-arreseted cells. A) HeLa cells were transfected with Nuf2-mCherry, PP1-mCherry or Nuf2PP1-mCherry. mCherry levels were quantified. Nuf2PP1 expression was lower than Nuf2 or PP1 expression. B) In cells treated with nocodazole, Ska3 at kinetochores was not different in cells expressing Nuf2-mCherry, PP1-mCherry, or Nuf2PP1-mCherry. C) Chromosomes were prepared from cells expressing Nuf2-mCherry, PP1-mCherry, or Nuf2PP1-mCherry. Cells expressing were arrested in mitosis with MG132. Western blotting (Figure 3H) reveals that cells expressing Nuf2-PP1 show higher levels of the APC/C component APC2 bound to mitotic chromosomes. D) HeLa cells transfected with Nuf2-mCherry, PP1-mCherry or Nuf2PP1-mCherry were imaged by time-lapse microscopy. Scatter plot shows mitotic progression is similar in cells expressing Nuf2-, PP1- or Nuf2PP1-mCherry.

